# AI-designed cyclic peptides enable controllable modulation of the CD28 immune checkpoint

**DOI:** 10.64898/2026.03.06.710051

**Authors:** Katarzyna Kuncewicz, Saurabh Upadhyay, Renjie Zhu, Hongliang Duan, Moustafa T. Gabr

## Abstract

Immune checkpoint therapies have transformed immunotherapy but remain dominated by biologic agents characterized by prolonged receptor occupancy and limited pharmacologic controllability. Synthetic modalities capable of targeting protein-protein interaction interfaces while enabling tunable immune regulation remain largely unexplored. Here, we report an AI-guided strategy for discovering cyclic peptide antagonists of the costimulatory receptor CD28. The lead peptide, CIP-3, binds the CD28 extracellular domain with nanomolar affinity and competitively disrupts CD28-ligand interactions. In primary human immune systems, CIP-3 suppresses CD28-dependent T-cell activation without intrinsic agonist activity and exhibits rapid pharmacologic reversibility, enabling exposure-dependent control of immune signaling. In a T-cell transfer model of chronic colitis, CIP-3 confers dose-dependent therapeutic efficacy and reduces systemic inflammatory cytokines. CIP-3 also suppresses cytokine production across independent healthy donors and patient-derived PBMCs from individuals with ulcerative colitis with efficacy comparable to a benchmark anti-CD28 biologic. Together, these findings establish AI-designed cyclic peptides as a controllable synthetic modality for immune checkpoint modulation.

## INTRODUCTION

Immune checkpoint pathways govern the magnitude, quality, and duration of adaptive immune responses and have emerged as central therapeutic targets across oncology, autoimmune disease, and transplantation medicine (1–3). Therapeutic blockade of inhibitory checkpoints such as PD-1 and CTLA-4 has transformed clinical practice and validated immune modulation as a powerful therapeutic strategy (4–6). Increasing attention has now turned toward costimulatory pathways that directly regulate T-cell activation thresholds and effector function (7). Among these, the receptor CD28 plays a central role in amplifying T-cell receptor signaling through interactions with its ligands CD80 and CD86 on antigen-presenting cells, promoting cytokine production, proliferation, metabolic reprogramming, and survival of T lymphocytes (8–10). Dysregulated CD28 signaling contributes to inflammatory and autoimmune diseases, including rheumatoid arthritis and inflammatory bowel disease (IBD), motivating efforts to therapeutically modulate this pathway (11–13).

To date, immune checkpoint modulation has been dominated by biologic agents, including monoclonal antibodies and receptor fusion proteins, which effectively engage extended protein-protein interaction surfaces (14–16). Despite their success, biologics present intrinsic limitations including prolonged receptor occupancy, limited pharmacologic controllability, immunogenicity risk, and complex manufacturing requirements (17–19). Moreover, modulation of costimulatory receptors such as CD28 carries unique safety considerations, as excessive receptor activation can trigger severe cytokine release syndromes (20). These limitations highlight the need for therapeutic strategies capable of precisely tuning immune signaling while maintaining high specificity for receptor interfaces.

Developing synthetic modulators of immune checkpoint receptors remains challenging because their extracellular domains typically present shallow, solvent-exposed protein-protein interaction surfaces that lack the deep binding pockets commonly exploited by small molecule drugs (21,22). Consequently, most successful therapeutic approaches rely on large biologic agents capable of engaging extended receptor surfaces, leaving a substantial modality gap between conventional small molecules and antibodies (23).

Cyclic peptides represent an emerging molecular class capable of bridging this gap. Their conformationally constrained architectures enable recognition of extended protein surfaces with greater specificity than small molecules while retaining synthetic accessibility, modular optimization potential, and tunable pharmacologic properties distinct from biologics (24–27). Cyclic scaffolds have demonstrated the ability to engage challenging protein-protein interfaces across diverse biological systems, including intracellular signaling proteins and extracellular receptors (28,29). However, systematic strategies for discovering cyclic peptide modulators of immune checkpoints remain limited, in part because efficient exploration of peptide sequence and conformational space poses significant design challenges.

Recent advances in AI-guided molecular design provide new opportunities to address these challenges. Machine-learning approaches integrating structural prediction, sequence optimization, and interaction modeling are increasingly enabling rational discovery of ligands targeting complex protein interfaces previously considered difficult to drug (30–33). Applying such strategies to immune checkpoint receptors could enable the development of synthetic modulators with improved pharmacologic controllability.

Here we sought to establish cyclic peptides as a controllable synthetic modality for targeting immune checkpoint receptors. Using an AI-guided discovery strategy, we identified cyclic peptide antagonists targeting the CD28 immune checkpoint and demonstrated that these molecules enable tunable modulation of T-cell costimulatory signaling. AI-designed cyclic peptides achieve nanomolar binding to CD28, disrupt receptor–ligand interactions, and suppress T-cell activation across multiple human immune systems. Importantly, peptide-mediated antagonism exhibits reversible pharmacology and absence of intrinsic agonist activity, distinguishing it from antibody-based approaches. Cross-reactivity with murine CD28 and pharmacokinetic (PK) characterization enabled in vivo evaluation, where cyclic peptide treatment produced dose-dependent therapeutic efficacy in a T-cell transfer model of chronic colitis. Together, these findings establish cyclic peptides as a synthetic modality capable of controllable immune checkpoint modulation and provide a framework for next-generation immunotherapies targeting costimulatory receptors.

## RESULTS

### AI-guided discovery identifies cyclic peptide ligands targeting the CD28 extracellular interface

To enable synthetic modulation of the CD28 costimulatory receptor, we developed an AI-guided discovery framework integrating reinforcement learning-based sequence optimization with structure prediction to generate cyclic peptides predicted to engage the extracellular domain of CD28 (Fig. 1A). CD28 presents a relatively shallow and solvent-exposed protein-protein interaction surface, a structural topology that has historically limited the applicability of conventional small molecule approaches. In contrast, short peptides can present extended contact surfaces that more closely approximate native ligand geometries, rendering them well suited for targeting interfaces governed by geometry-dependent extracellular recognition rather than enzymatic activity. Disulfide-constrained cyclic peptides combine compact size with conformational preorganization, enabling stable presentation of interaction motifs across flat or discontinuous protein surfaces while reducing entropic penalties upon binding and improving resistance to proteolytic degradation.

**Figure 1.**
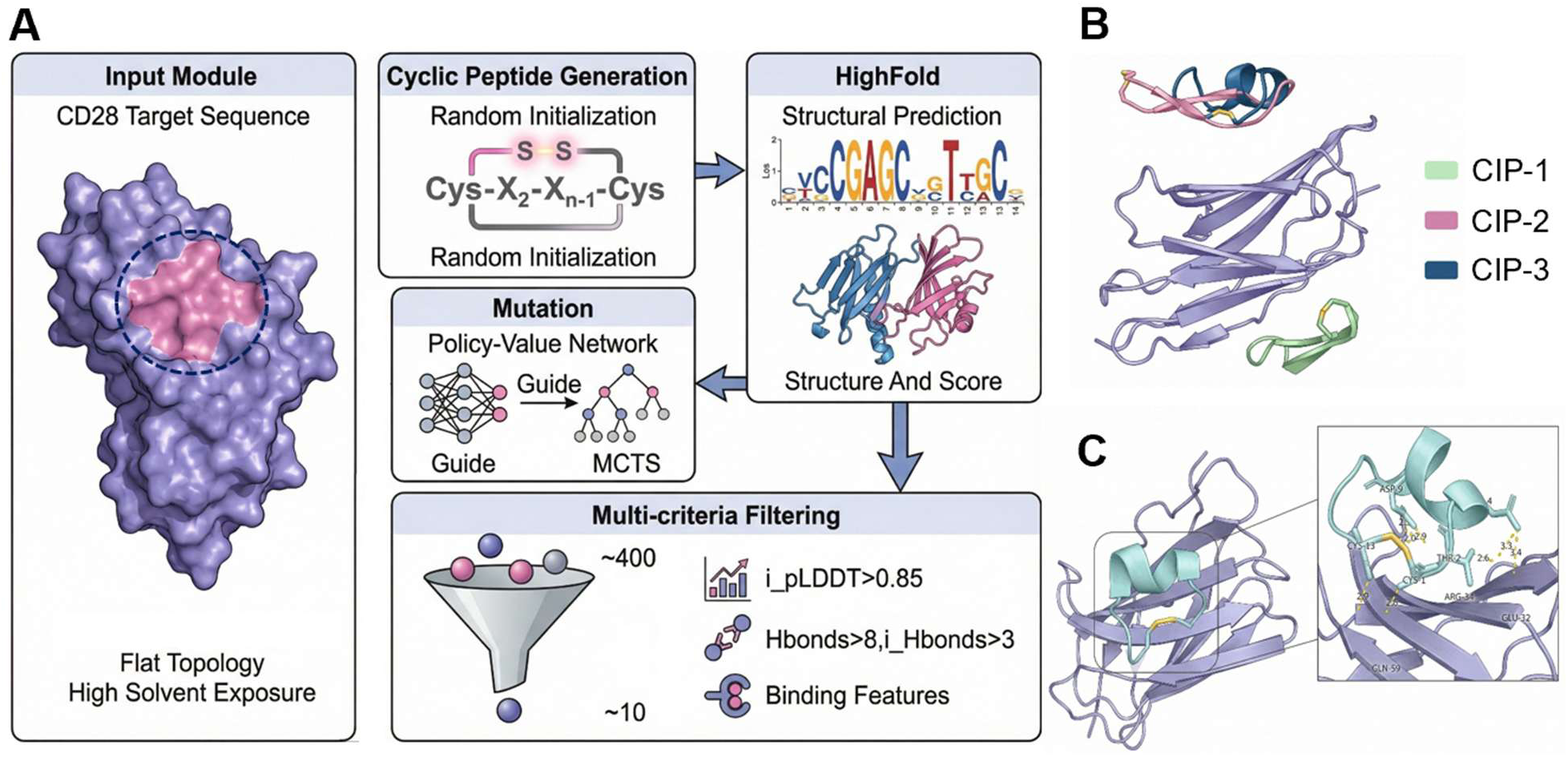
AI-guided design and structural characterization of cyclic immune peptides targeting CD28. **(A)** Computational workflow for cyclic peptide discovery. A solvent-exposed, topologically flat region of CD28 was selected as the target interface and used to guide de novo cyclic peptide generation with a disulfide-constrained scaffold (Cys-X_2_-X_n-1_-Cys). Candidate sequences were iteratively optimized using a policy-value network with Monte Carlo tree search (MCTS) and evaluated by HighFold structural prediction. Multi-criteria filtering based on predicted structural confidence (i_pLDDT > 0.85), intermolecular hydrogen bonding, and binding features reduced ∼400 candidates to ∼10 high-confidence leads. **(B)** Structural comparison of the designed cyclic peptides (CIP-1, CIP-2, and CIP-3) bound to CD28. The peptides occupy the targeted interface region identified in panel A. **(C)** Predicted binding mode of the lead peptide (CIP-3) in complex with CD28, with inset highlighting key intermolecular interactions and hydrogen bonds stabilizing the interface.

We therefore prioritized conformationally constrained macrocyclic scaffolds capable of achieving high surface complementarity to the CD28 interface. Using the Highplay platform (34), which integrates Monte Carlo Tree Search-based reinforcement learning with the HighFold structure prediction model (35), candidate cyclic peptide sequences were generated directly from the CD28 amino-acid sequence and ranked using predicted interaction energies, interface hydrogen-bonding potential, and structural confidence metrics derived from ensemble modeling. Filtering criteria included high predicted interface accuracy (i_pLDDT > 0.85) together with favorable intermolecular interaction features, enabling efficient exploration of chemically tractable binders targeting a structurally challenging receptor surface.

Three disulfide-constrained cyclic peptides were selected for experimental evaluation. We refer to these molecules as cyclic immune peptides (CIPs), reflecting both their cyclic architecture and intended immune-modulatory function. Structural modeling predicted that the peptides occupy regions proximal to the CD28 ligand-binding interface responsible for CD80/CD86 engagement, consistent with a mechanism of competitive antagonism of receptor-ligand interactions (Fig. 1B,C; fig. S1,2 for additional models).

### Cyclic peptide CIP-3 binds CD28 with nanomolar affinity

Binding interactions between the designed cyclic peptides and the human CD28 extracellular domain (ECD) were quantified using microscale thermophoresis (MST) under solution-phase conditions. This approach enables sensitive detection of molecular interactions without immobilization, an important consideration for targets such as CD28 that present relatively shallow, protein-protein interaction-driven binding surfaces.

Among the three candidates, CIP-1 did not exhibit detectable binding across the tested concentration range, whereas CIP-2 displayed concentration-dependent interaction with an apparent dissociation constant in the micromolar range (Kd = 47.16 μM, 95% CI: 28-164 μM, Fig. 2B). In contrast, CIP-3 demonstrated substantially stronger engagement, yielding a well-defined sigmoidal binding curve with a Kd value of approximately 108 nM, 95% CI: 77-144 nM (Fig. 2C). The nearly 400-fold affinity difference between CIP-2 and CIP-3 underscores the critical role of sequence-dependent structural features in governing productive CD28 recognition, despite the shared disulfide-constrained cyclic architecture.

**Figure 2.**
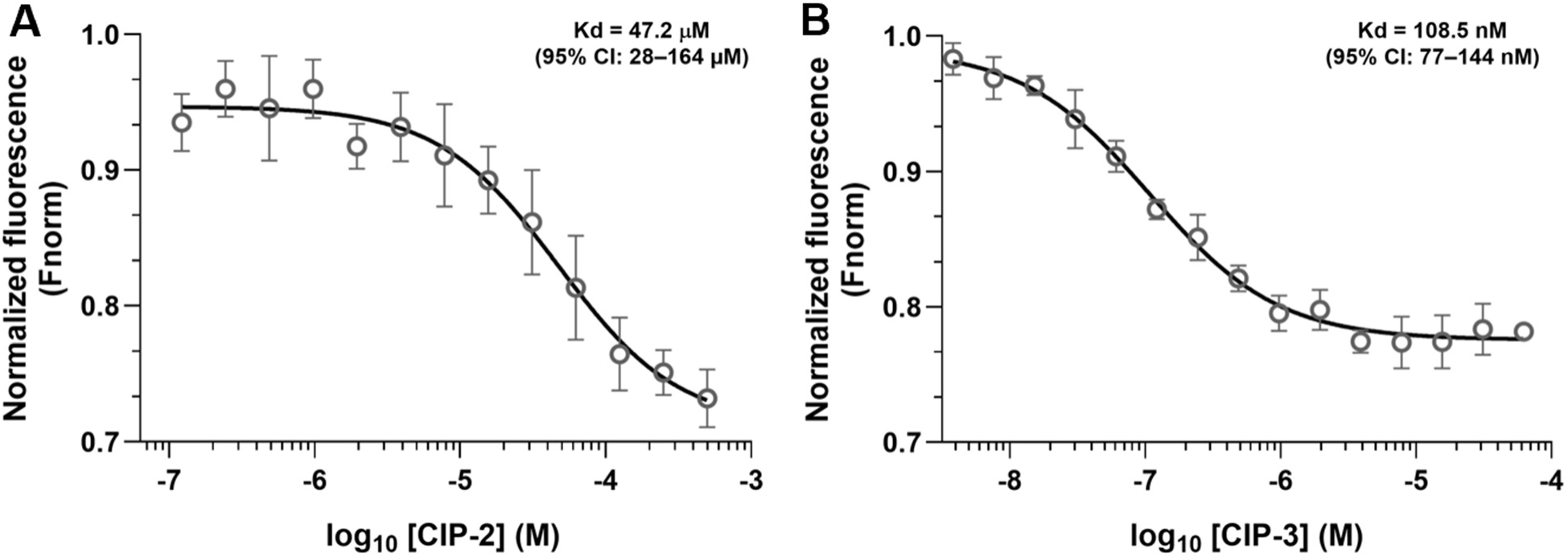
Microscale thermophoresis analysis of cyclic peptide binding to CD28. Binding of disulfide-constrained cyclic peptides to the human CD28 extracellular domain (ECD) was quantified by microscale thermophoresis (MST) using intrinsic fluorescence detection (spectral shift mode; Monolith X). Normalized fluorescence (Fnorm, 670/650 nm) is plotted as a function of peptide concentration. **(A)** CIP-2 exhibits concentration-dependent binding consistent with a moderate-affinity interaction. **(B)** CIP-3 displays a well-defined sigmoidal binding curve indicative of higher-affinity engagement with CD28. Data points represent mean ± SD from three independent measurements. Binding curves were fitted using nonlinear regression assuming a 1:1 binding model to estimate equilibrium dissociation constants (Kd).

The absence of detectable binding for CIP-1 provides an internal specificity control, arguing against nonspecific interactions driven solely by peptide cyclization or physicochemical properties. Instead, the observed affinity hierarchy supports a model in which effective CD28 engagement requires precise sequence-encoded surface complementarity combined with conformational preorganization imposed by cyclization. Collectively, these findings demonstrate that compact, disulfide-constrained peptides can achieve nanomolar binding affinity to a costimulatory receptor characterized by a shallow extracellular interface.

### CIP-3 disrupts CD28-ligand interactions through competitive antagonism

The ability of CD28-binding cyclic peptides to functionally disrupt receptor-ligand engagement was evaluated using a competitive binding assay. Consistent with biophysical measurements, both CIP-2 and CIP-3 inhibited CD28-CD80 interactions in a concentration-dependent manner, yielding well-defined sigmoidal inhibition curves (Fig. 3A,B). CIP-2 displayed moderate inhibitory activity (IC_50_ = 59.3 µM, 95% CI: 34-242 μM), whereas CIP-3 exhibited substantially greater potency (IC_50_ = 609 nM, 95% CI: 419-874 nM), representing an approximately 65-fold improvement in functional activity.

**Figure 3.**
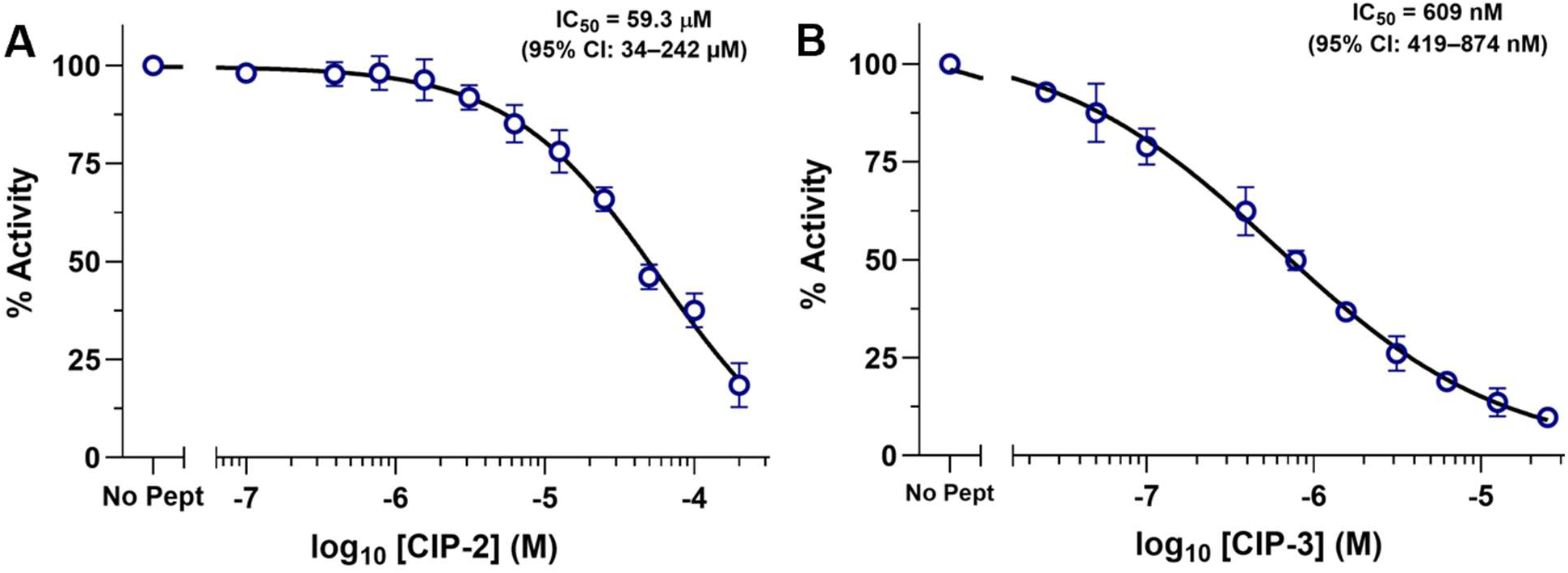
Cyclic peptides competitively disrupt CD28-CD80 interactions. Functional inhibition of CD28-ligand engagement by cyclic peptides was assessed using a competitive binding assay in which peptide-mediated blockade of biotinylated CD80 binding to immobilized recombinant human CD28 was quantified. **(A)** CIP-2 inhibits CD28-CD80 interaction in a concentration-dependent manner with moderate potency. **(B)** CIP-3 exhibits substantially enhanced inhibitory activity, yielding a well-defined sigmoidal inhibition curve with submicromolar potency. Data represent mean ± SD from three independent experiments. IC_50_ values were determined by nonlinear regression analysis.

The close agreement between MST-derived binding affinities and functional inhibition supports a mechanism in which cyclic peptides directly occupy the CD28 ligand-binding interface rather than producing nonspecific assay interference. Notably, the submicromolar potency of CIP-3 demonstrates that a compact disulfide-constrained peptide can effectively compete with native costimulatory interactions at an immune checkpoint receptor, a property traditionally achieved using biologic agents. Having established that cyclic peptides disrupt CD28-CD80 interactions in a cell-free system, we next evaluated whether this antagonism translates into modulation of CD28-dependent signaling in cellular assays.

### Cyclic peptide antagonists suppress CD28-dependent signaling in cellular systems

The functional consequences of CD28 antagonism were evaluated using a luciferase reporter assay that quantitatively measures CD28 costimulatory signaling in Jurkat T cells co-cultured with antigen-presenting cells. Peptides capable of binding CD28 and disrupting CD28-CD80 interactions suppressed reporter activity in a concentration-dependent manner (Fig. 4). CIP-2 produced moderate inhibition (IC_50_ = 10.1 µM, 95% CI: 6.3-30.5 μM, Fig. 4A), whereas CIP-3 exhibited markedly enhanced potency with a submicromolar IC_50_ of 443 nM, 95% CI: 350-534 nM (Fig. 4B).

**Figure 4.**
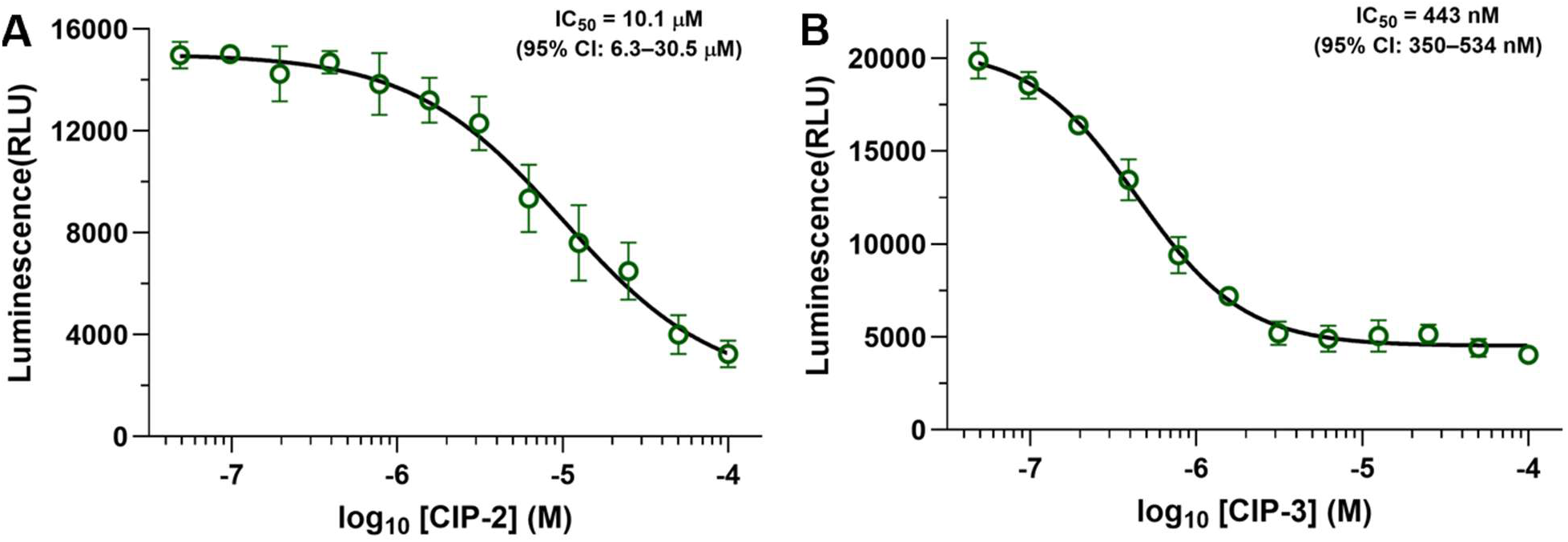
Cyclic peptides inhibit CD28-dependent costimulatory signaling in cells. Functional antagonism of CD28 signaling was evaluated using a reporter assay in which Jurkat T cells were co-cultured with antigen-presenting cells to induce CD28-dependent activation, quantified by an NFAT-driven luciferase readout. **(A)** CIP-2 inhibits CD28 signaling in a concentration-dependent manner with moderate potency. **(B)** CIP-3 exhibits substantially enhanced inhibitory activity, achieving submicromolar inhibition. Data represent mean ± SEM from at least three independent experiments. Concentration-response curves were fitted using a four-parameter logistic regression model.

The rank order of cellular activity closely mirrored trends observed in MST binding and ELISA-based competition assays, supporting a direct relationship between peptide engagement of the CD28 extracellular domain and downstream signaling blockade. These findings demonstrate that disulfide-constrained cyclic peptides can functionally antagonize CD28 signaling in a cellular environment, establishing CIP-3 as a lead peptide with robust activity.

### CIP-3 reproduces biologic-like immune modulation in primary human systems

To evaluate whether CIP-3 could functionally modulate CD28-mediated T-cell activation in primary human systems, peripheral blood mononuclear cells (PBMCs) from independent healthy donors (n = 10 donors) were stimulated with anti-CD3/CD28 antibodies in the presence of increasing concentrations of the cyclic peptide (Fig. 5A). CIP-3 produced a clear concentration-dependent suppression of cytokine secretion, with progressive reductions observed in both IL-2 and IFN-γ levels (Fig. 5B-C). Importantly, inhibitory activity was reproducible across a cohort of ten independent donors, despite expected inter-individual variability in baseline cytokine responses, supporting a consistent pharmacologic effect across heterogeneous immune backgrounds.

**Figure 5.**
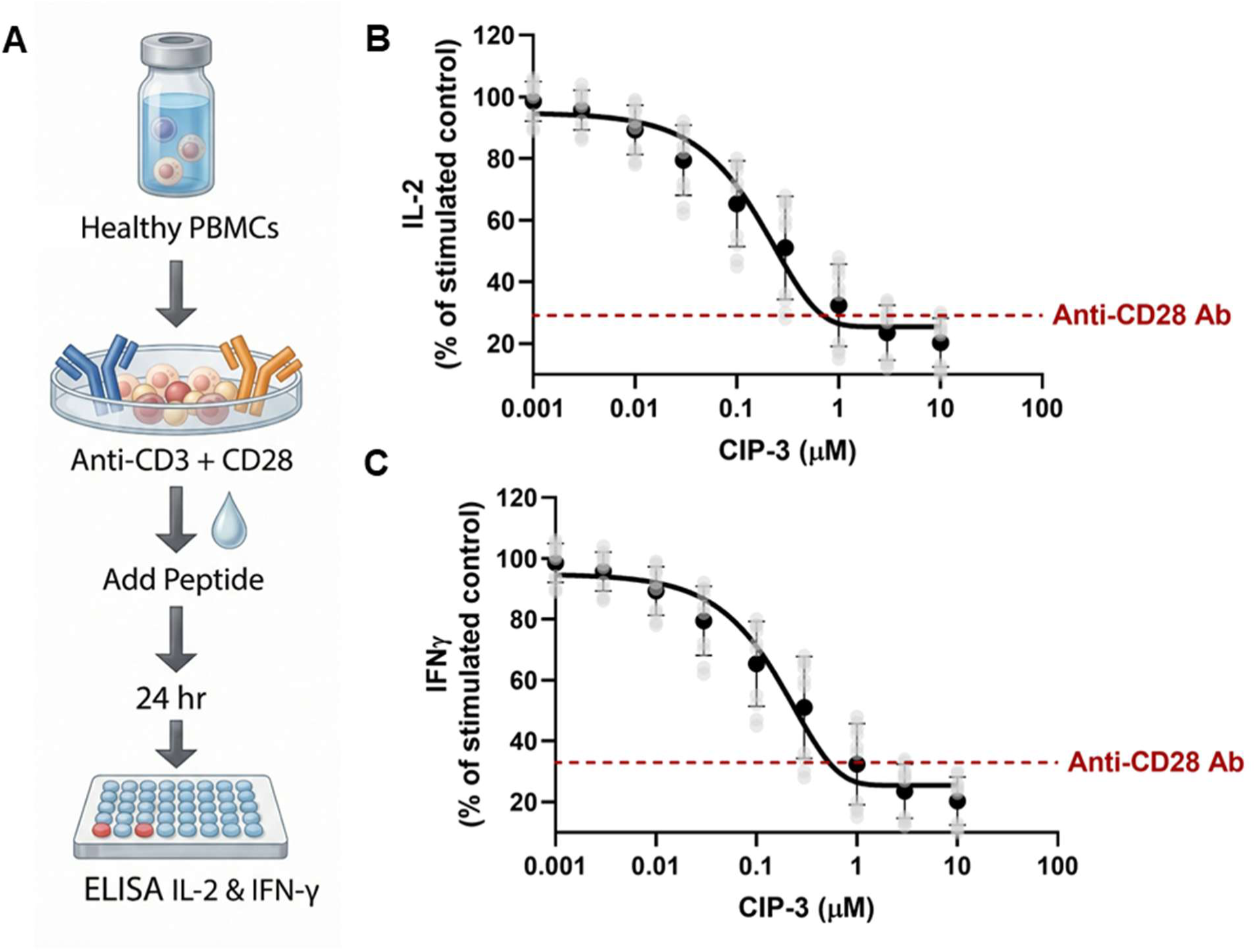
Cyclic CD28 peptide suppresses primary human T-cell activation across independent donors. **(A)** Experimental schematic showing stimulation of peripheral blood mononuclear cells (PBMCs) from healthy donors with anti-CD3/CD28 antibodies followed by treatment with cyclic peptide CIP-3 and cytokine measurement after 24 h. **(B-C)** Concentration-dependent inhibition of IL-2 **(B)** and IFN-γ **(C)** secretion by CIP-3 across independent donors (n = 10). Light gray symbols represent individual donor responses, and black symbols indicate mean ± SEM. A benchmark anti-CD28 blocking antibody (FR104, 10 µg mL^−1^) is shown for comparison (red dashed line).

At higher peptide concentrations, the magnitude of cytokine suppression approached that achieved with a benchmark anti-CD28 blocking antibody (FR104) under matched experimental conditions, indicating that the cyclic peptide can produce biologically meaningful attenuation of CD28 co-stimulatory signaling. Nonlinear regression analysis of the concentration-response relationship revealed sub-micromolar functional potency, further supporting effective engagement of the CD28 signaling axis.

### CIP-3 exhibits no intrinsic agonist activity and mediates reversible checkpoint modulation

Given historical safety concerns associated with CD28 targeting, we first evaluated whether CIP-3 alone could induce immune activation in the absence of co-stimulatory signaling. Treatment of primary PBMCs with CIP-3 across the tested concentration range (0.01-10 µM) did not increase cytokine production relative to unstimulated baseline controls (Fig. 6A,B), indicating absence of intrinsic agonist activity. These findings suggest that receptor engagement by the cyclic peptide does not trigger unintended CD28 signaling and supports a favorable functional safety profile compared with agonistic CD28 antibodies.

**Figure 6.**
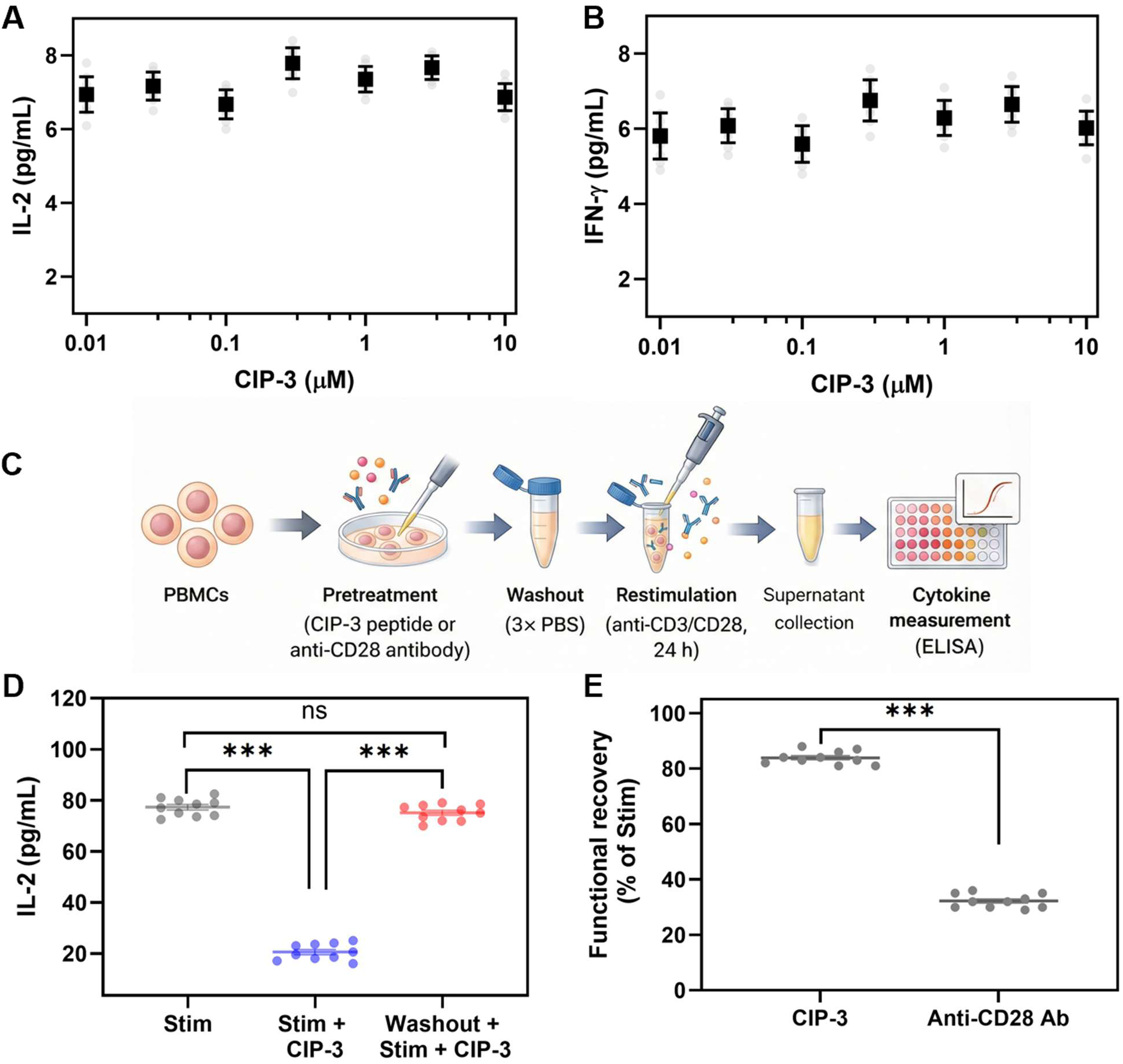
CIP-3 lacks intrinsic agonist activity and mediates reversible modulation of CD28 signaling. **(A,B)** Primary PBMCs treated with CIP-3 (0.01-10 µM) in the absence of stimulation show no increase in cytokine production relative to unstimulated controls, indicating absence of intrinsic agonist activity. IL-2 **(A)** and IFN-γ **(B)** concentrations were measured by ELISA. **(C)** Experimental schematic illustrating pretreatment with CIP-3 or anti-CD28 antibody, washout (3× PBS), and subsequent restimulation with anti-CD3/CD28 prior to cytokine measurement. **(D)** Functional recovery following inhibitor removal demonstrates rapid reversibility of peptide-mediated inhibition (CIP-3, 1 μM) compared with sustained inhibition following anti-CD28 antibody treatment (10 µg/mL). **(E)** Quantification of functional recovery expressed as percentage of stimulated control highlights substantially greater recovery following CIP-3 washout relative to antibody treatment. Data represent mean ± SEM from independent donors (n = 10). Statistical analysis was performed using one-way ANOVA with multiple comparisons unless otherwise indicated. ns, not significant; \*\*\**P* < 0.001.

We next examined whether peptide-mediated inhibition could be pharmacologically controlled through reversible receptor engagement. In washout experiments, PBMCs were preincubated with CIP-3 (1 µM) or a benchmark anti-CD28 antibody (1 µg mL^−1^), followed by removal of unbound inhibitor prior to stimulation (Fig. 6C). Peptide-treated cells exhibited rapid recovery of signaling activity, reaching approximately 85% of stimulated control levels following washout, whereas biologic-mediated inhibition persisted with only approximately 30% recovery over the same period (Fig. 6D,E). These results demonstrate that CIP-3 engages CD28 in a transient and reversible manner, consistent with noncovalent ligand-receptor interactions, in contrast to the prolonged receptor occupancy associated with antibody binding.

The rapid reversibility observed with CIP-3 indicates that synthetic cyclic ligands can enable tunable modulation of CD28 signaling with temporal control distinct from antibody-based approaches. Importantly, the combination of absent intrinsic agonism and controllable pharmacological reversibility suggests a therapeutic modality with a potentially improved safety window compared with conventional biologic checkpoint modulators. Results were reproducible across independent donors (n = 10), supporting robustness across heterogeneous immune backgrounds.

### CIP-3 suppresses CD28-mediated cytokine production in T cells derived from patients with ulcerative colitis and demonstrates efficacy comparable to a benchmark biologic

To determine whether cyclic peptides can achieve functionally meaningful modulation of T-cell co-stimulation in disease-relevant human systems, we evaluated CIP-3 using PBMCs obtained from patients with ulcerative colitis (n = 5). Upon anti-CD3/CD28 stimulation, CIP-3 induced a concentration-dependent attenuation of cytokine production, suppressing both IL-2 and IFN-γ secretion across donors (Fig. 7A,B). Notably, inhibition plateaued at micromolar concentrations, consistent with saturable but incomplete pathway modulation rather than binary receptor blockade.

**Figure 7.**
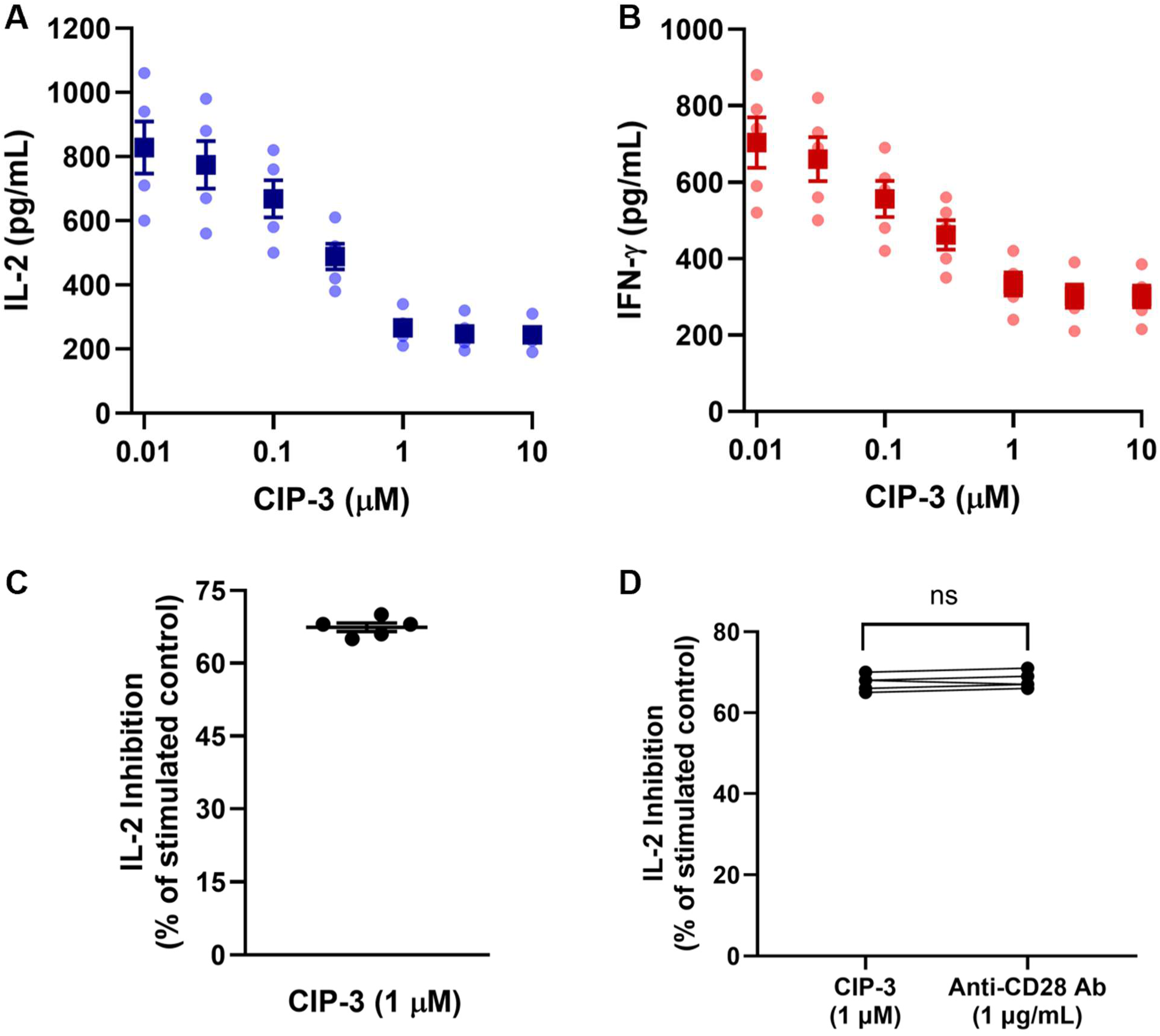
CIP-3 suppresses CD28-mediated cytokine production in PBMCs from patients with ulcerative colitis with efficacy comparable to a benchmark biologic. **(A,B)** PBMCs obtained from donors with ulcerative colitis (n = 5) were stimulated with anti-CD3/CD28 and treated with increasing concentrations of CIP-3 (0.01-10 µM). CIP-3 produced dose-dependent inhibition of IL-2 **(A)** and IFN-γ **(B)** secretion measured by ELISA. Data points represent individual donors with mean ± SEM. **(C)** IL-2 inhibition across independent patient samples at 1 µM CIP-3 demonstrates consistent activity in disease-derived primary immune cells. **(D)** Comparison of peptide-mediated and antibody-mediated inhibition across matched patient samples shows that CIP-3 (1 µM) achieves levels of IL-2 inhibition comparable to those obtained with an anti-CD28 monoclonal antibody (10 µg/mL) (paired two-tailed t-test, ns).

This graded suppression profile contrasts with the fixed amplitude typically associated with monoclonal antibody-mediated inhibition and suggests that peptide engagement permits pharmacologic tuning of co-stimulatory signaling intensity. Across independent patient samples, CIP-3 reproducibly attenuated IL-2 production at a pharmacologically relevant concentration (1 µM), with limited inter-donor variability (Fig. 7C), indicating robust activity within heterogeneous inflammatory immune environments.

To benchmark functional efficacy, we directly compared peptide-mediated modulation with a clinically relevant anti-CD28 monoclonal antibody (FR104) in matched patient samples. CIP-3 achieved levels of IL-2 suppression comparable to antibody treatment, with no statistically significant difference between conditions (Fig. 7D). Collectively, these results establish that CD28 co-stimulatory signaling can be modulated in a tunable, pharmacologically controllable manner using a non-antibody ligand, supporting the feasibility of peptide-based immune checkpoint modulation in inflammatory disease.

### CIP-3 demonstrates dose-dependent therapeutic efficacy in a T-cell transfer model of colitis

To evaluate the in vivo therapeutic potential of CIP-3, we employed the adoptive CD4⁺CD45RB^high^ T-cell transfer model of chronic colitis in C.B-17 scid mice (Fig. 8A). This model recapitulates key features of human IBD, including progressive weight loss, diarrhea, colonic shortening, and elevated systemic inflammatory cytokines (36–38). We first established functional cross-reactivity with murine CD28 to ensure translational relevance of the in vivo model. In primary murine splenocytes stimulated with anti-CD3/CD28, CIP-3 dose-dependently suppressed T-cell activation and cytokine production, confirming engagement of murine CD28 signaling (fig. S6).

**Figure 8.**
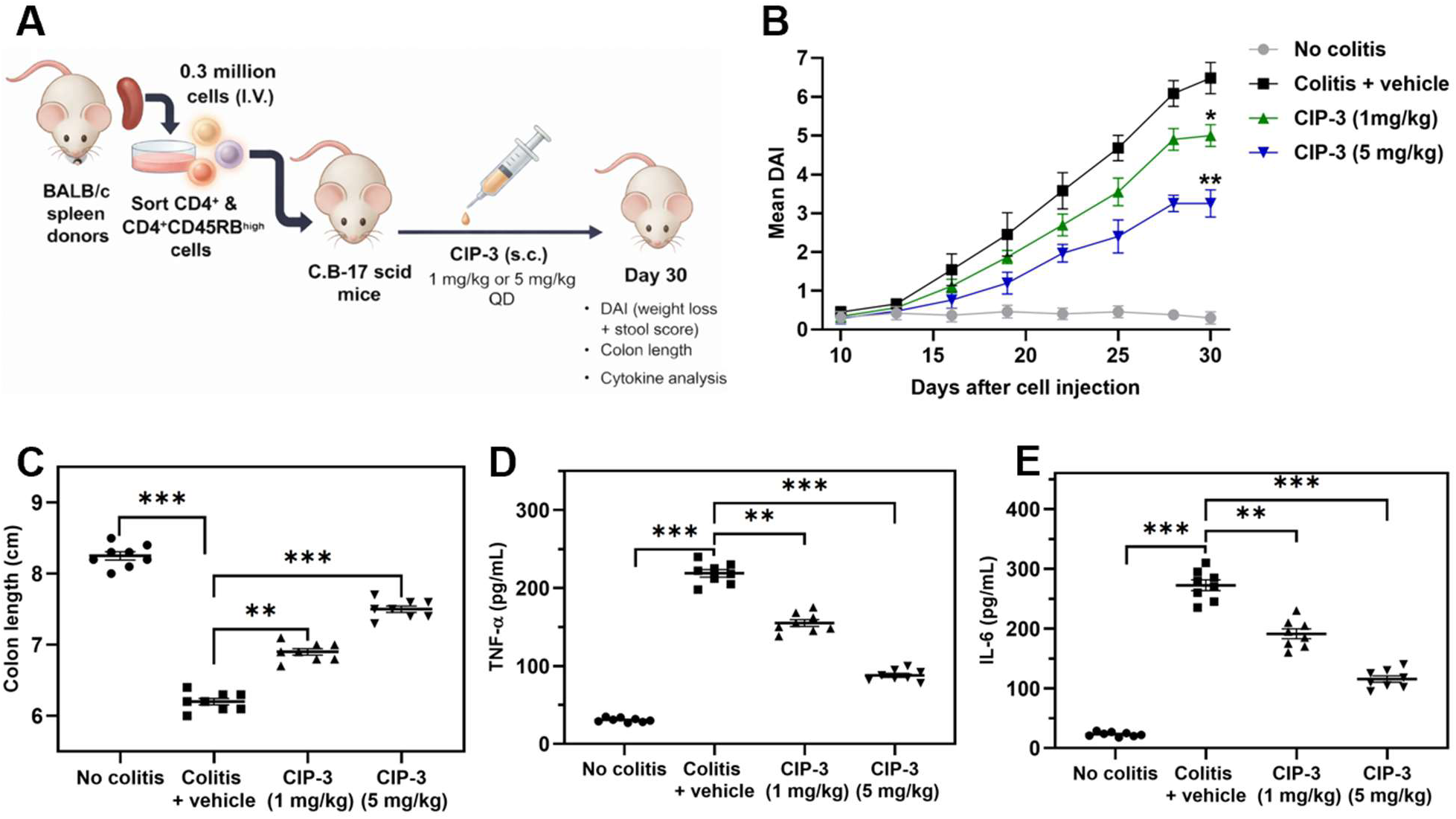
CIP-3 confers dose-dependent therapeutic efficacy in a T-cell transfer model of colitis. **(A)** Schematic representation of the adoptive CD4⁺CD45RB^high^ T-cell transfer model of chronic colitis. CD4⁺CD45RB^high^ T cells were isolated from BALB/c donor spleens and intravenously transferred into C.B-17 scid recipient mice. Beginning after cell transfer, mice received daily subcutaneous administration of CIP-3 (1 mg/kg or 5 mg/kg) or vehicle. Disease progression was monitored for 30 days. Endpoints included disease activity index (DAI), colon length, and serum cytokine analysis. **(B)** CIP-3 reduced disease severity in a dose-dependent manner. DAI (combined weight loss and stool score) was assessed longitudinally. Vehicle-treated mice developed progressive colitis, whereas CIP-3 treatment significantly attenuated disease progression, with greater efficacy observed at 5 mg/kg. **(C)** CIP-3 mitigated colonic shortening associated with inflammation. Colon length was measured on day 30. Vehicle-treated mice exhibited marked shortening relative to no-colitis controls, whereas CIP-3 treatment significantly restored colon length in a dose-dependent manner. **(D-E)** CIP-3 suppressed systemic inflammatory cytokines. Serum TNF-α (**D)** and IL-6 **(E)** levels were quantified by ELISA at study termination. Vehicle-treated mice showed elevated cytokine concentrations, while CIP-3 significantly reduced TNF-α and IL-6 levels, with stronger suppression at 5 mg/kg. Data represent mean ± SEM (n = 8 per group). Statistical analysis was performed using one-way ANOVA with Tukey’s multiple comparison test for endpoint analyses and two-way ANOVA for longitudinal DAI measurements. \**P* < 0.05, \*\**P* < 0.01, \*\*\**P < 0.001* versus colitis + vehicle group.

Prior to in vivo efficacy studies, we characterized the pharmacokinetic (PK) and metabolic stability profile of CIP-3. In mouse plasma, CIP-3 demonstrated high stability, with >85% parent compound remaining after 6 h at 37 °C. In liver microsomes, CIP-3 exhibited low intrinsic clearance in both mouse (Cl_int_ = 6.8 µL/min/mg protein) and human (Cl_int_ = 4.9 µL/min/mg protein) systems, corresponding to microsomal half-lives of 102 min (mouse) and 138 min (human), respectively. Following a single subcutaneous administration in C57BL/6 mice, CIP-3 displayed: C_max_ (5 mg/kg): 3.4 ± 0.6 µg/mL, C_max_ (1 mg/kg): 0.72 ± 0.11 µg/mL, T_max_: 0.5-1 h, Terminal half-life (t_1/2_): 3.9 ± 0.4 h, apparent clearance (CL/F): 0.42 L/h/kg, and AUC_0-24h_ (5 mg/kg): 12.8 ± 1.9 µg·h/mL. Importantly, plasma concentrations remained above the in vitro functional concentrations for CD28 inhibition (≈ 500 nM) for approximately 10-12 h at 5 mg/kg and 4-6 h at 1 mg/kg. These exposure levels and half-life parameters supported a once-daily subcutaneous dosing regimen and justified evaluation of 1 mg/kg and 5 mg/kg in the T-cell transfer colitis model.

Vehicle-treated mice developed progressive colitis characterized by increasing disease activity index (DAI) scores over 30 days (Fig. 8B). In contrast, CIP-3 significantly reduced disease severity in a dose-dependent manner. Mice treated with 1 mg/kg CIP-3 exhibited partial attenuation of DAI progression, whereas 5 mg/kg treatment produced a more pronounced reduction in disease burden, demonstrating clear pharmacodynamic dose responsiveness. Chronic inflammation in vehicle-treated mice resulted in marked colon shortening at study termination (Fig. 8C). Treatment with CIP-3 significantly restored colon length relative to vehicle controls. The 1 mg/kg group showed partial structural rescue, while 5 mg/kg treatment more robustly preserved colon length, consistent with the observed improvements in clinical scores.

To assess the impact of CIP-3 on inflammatory mediators, serum cytokine levels were quantified at day 30. Vehicle-treated colitic mice displayed elevated TNF-α and IL-6 concentrations (Fig. 8D-E). CIP-3 significantly reduced both cytokines in a dose-dependent manner, with greater suppression observed at 5 mg/kg. These findings indicate that CIP-3 dampens pathogenic T-cell-driven inflammatory responses in vivo. Together, these data demonstrate that CIP-3 confers dose-dependent therapeutic benefit in a T-cell-mediated model of colitis, with concordant improvements in clinical disease indices, anatomical preservation of colon length, and systemic cytokine suppression. The in vivo efficacy is supported by validated murine CD28 engagement and PK properties that sustain target-relevant exposure under once-daily subcutaneous administration.

## DISCUSSION

The present study establishes cyclic peptides as a controllable synthetic modality for targeting immune costimulatory receptors traditionally addressed using biologic agents. Through an AI-guided discovery strategy, we identified a disulfide-constrained cyclic peptide capable of nanomolar binding to the extracellular domain of CD28 and demonstrated that this molecule achieves biologic-like functional suppression of T-cell activation across primary human immune systems. Importantly, this synthetic ligand exhibits reversible pharmacology, absence of intrinsic agonist activity, and dose-dependent therapeutic efficacy in vivo. These findings provide proof of principle that compact cyclic scaffolds can function as systemically active modulators of immune checkpoint signaling.

Targeting costimulatory receptors with synthetic ligands has historically been constrained by structural considerations. The extracellular domains of receptors such as CD28 present shallow protein-protein interaction surfaces that lack the deep binding pockets typically exploited by small molecules. Our results demonstrate that conformationally constrained cyclic peptides can effectively engage these interfaces with nanomolar affinity and competitively disrupt receptor-ligand interactions. By presenting preorganized interaction motifs across extended surfaces, cyclic peptides can achieve high surface complementarity while retaining the synthetic tractability and modular optimization potential of chemically defined ligands.

A defining feature of this approach is pharmacologic controllability. In contrast to antibody-based checkpoint blockade, which is often characterized by prolonged receptor occupancy and sustained signaling modulation, peptide-mediated antagonism of CD28 was rapidly reversible following washout. The absence of intrinsic agonist activity further distinguishes this synthetic ligand from antibody-based strategies that carry risk of unintended receptor activation. Together, these properties suggest that cyclic peptide antagonists may enable exposure-dependent tuning of immune signaling intensity, providing temporal control over immune modulation.

The translational relevance of this modality is supported by activity across multiple biological systems. CIP-3 reproducibly suppressed cytokine production in PBMCs derived from independent healthy donors and from patients with ulcerative colitis, demonstrating functional robustness across heterogeneous immune backgrounds. PK characterization revealed favorable plasma stability and sustained systemic exposure following subcutaneous administration, enabling target-relevant engagement in vivo. In a T-cell transfer model of chronic colitis, cyclic peptide treatment produced dose-dependent reductions in disease activity, preservation of colon length, and suppression of systemic inflammatory cytokines, establishing proof of concept that synthetic checkpoint modulation can translate into therapeutic benefit.

Several considerations warrant further investigation. Although the present study demonstrates efficacy in a T-cell-mediated inflammatory model, additional work will be required to evaluate long-term safety, dosing flexibility, and activity across other disease contexts. Further optimization of peptide stability, formulation, and delivery strategies may enhance PK performance and expand therapeutic applications.

In summary, these findings establish AI-designed cyclic peptides as a viable synthetic modality for immune checkpoint modulation. By combining structural complementarity to protein–protein interaction interfaces with exposure-dependent pharmacologic control, cyclic peptides provide a bridge between small molecules and biologics and offer a foundation for next-generation immunotherapies targeting costimulatory receptors.

## MATERIALS AND METHODS

### AI-guided cyclic peptide design

A series of data preprocessing steps were performed prior to model-based design to ensure the compatibility of Protein Data Bank (PDB) files with our computational workflow. The design of CD28 binders was based on the PDB structure 1YJD; the amino acid sequence of chain C was extracted using a custom Python script and subsequently used as the input sequence of the CD28 target for the Highplay platform (34).

The binding epitope of CD28 consists of a relatively flat β-sheet domain. Given our focus on the interfacial binding characteristics of peptide-target complexes, a threshold of interface predicted local distance difference test (i_pLDDT) > 0.85 was applied in this study to ensure the structural accuracy of the peptide-CD28 binding interface. On the premise of validating the structural rationality of the binding interface, further screening was conducted with the criteria of number of hydrogen bonds (Hbonds) > 8 in the whole complex and interface hydrogen bonds (i_Hbonds) > 3 at the peptide-target contact site. These criteria were set to guarantee sufficient side chain-mediated intermolecular interactions between cyclic peptides and the CD28 surface, thus identifying candidate molecules with high binding affinity. Based on the aforementioned screening metrics, we established a standardized computational pipeline for the design and primary screening of CD28-targeting cyclic peptides. For the pre-screened cyclic peptide candidates, we performed in-depth structural analyses, including the characterization of side chain interaction modes with the CD28 target, the assessment of binding site accessibility, and a comprehensive evaluation of chemical synthetic feasibility. Ultimately, three cyclic peptide candidates, namely CIP-1 (CTAIHLDLRETEQC), CIP-2 (CSHVGFGRRQELLC), and CIP-3 (CTENWRRYDGPQC) were selected for experimental validation in vitro. All three candidates exhibited favorable computational metrics, displayed a high degree of structural complementarity to the CD28 binding interface, and thus possessed considerable potential for high CD28 binding affinity.

### Peptide synthesis and characterization

Peptide synthesis was carried out by solid phase peptide synthesis (SPPS) via the Fmoc/tBu strategy using an automated microwave peptide synthesizer - Liberty BLUE (CEM, Matthews, NC, USA). 424 mg Rink Amide ProTide Resin resin (0.59 mmol/g; CEM, Matthews, NC, USA) was used to synthesize each peptide. During the synthesis, each of the amino acids was coupled twice, using a 5-fold excess of the amount of resin deposition. The following solutions were used during the synthesis: OxymaPure/DIC as coupling reagents, 20% piperidine in DMF for Fmoc-deprotection, and DMF to wash between the deprotection and coupling steps. Synthesized peptides were cleaved from the resin using a mixture containing: 88% TFA, 5% H2O, 5% phenol and 2% TIPSI (10 ml of solution was used for 424 mg of resin). Crude peptides were precipitated with cold diethyl ether, decanted, and lyophilized. For purification, the peptides were dissolved in water with the addition of a 10-fold molar excess of dithiothreitol (DTT) over free sulfhydryl groups and incubated for 30 minutes at 60 °C. Peptides were purified by reversed phase HPLC, XBridge Prep C18 column (19 x 150 mm, 5 μm, Waters, MA, USA), flow rate 20 mL/min, 10 min run, 5-60% acetonitrile in water with 0.1% formic acid. The purity and mass spectra of the final products were analyzed using an ACQUITY HPLC system equipped with an SQ Detector 2 and an XBridge Prep C18 analytical column (4.6 × 150 mm, 5 μm, Waters, MA, USA), employing a linear gradient from 5% to 100% acetonitrile in water containing 0.1% formic acid over 10 minutes. Oxidation of the peptides was performed using compressed air. The peptide was dissolved in H2O and methanol (1:9, v:v), at a concentration of about 40 mg/L, and the pH was adjusted and kept between 8 and 9 using ammonia. The solution was stirred at room temperature for 7 days, and compressed air was run through the solution. After this time, the solvents were evaporated, and the peptides were lyophilized. Reaction progress was checked using analytical ACQUITY HPLC system equipped with an SQ Detector 2. After this process, the peptides were purified again using the same ACQUITY HPLC system on a XBridge Prep C18 column (19 x 150 mm, 5 μm, Waters, MA, USA). A linear gradient 5%-60% acetonitrile in water with 0.1% formic acid over 10 min was used.

### Recombinant proteins

Recombinant human CD28 extracellular domain (ECD) was obtained from Acro Biosystems (Catalog# CD8-H52H3) and reconstituted according to manufacturer instructions. Proteins were stored in aliquots at −80 °C to avoid repeated freeze-thaw cycles.

### Microscale thermophoresis (MST) binding assays

Binding affinities of cyclic peptides to the human CD28 ECD were determined by MST in spectral shift mode. His-tagged recombinant human CD28 ECD (Acro Biosystems) was fluorescently labeled using the RED-tris-NTA 2nd Generation dye (NanoTemper Technologies) according to the manufacturer’s protocol and diluted to a final concentration of 100 nM in assay buffer consisting of PBS (pH 7.0) supplemented with 0.005% Tween-20.

Cyclic peptides were prepared as serial dilutions in assay buffer and mixed 1:1 with labeled CD28 protein. Samples were incubated for 15 min at room temperature in the dark prior to measurement. MST experiments were performed at 25 °C using a Monolith X instrument (NanoTemper Technologies) and loaded into Monolith Capillaries.

Fluorescence was recorded at 670 nm and 650 nm, and normalized fluorescence (Fnorm) was calculated as the ratio F670/F650. Dissociation constants (Kd) were obtained by fitting concentration–response curves using a four-parameter nonlinear regression model in MO.Affinity Analysis software and GraphPad Prism 10. Reported values represent mean ± SD from at least three independent experiments.

### CD28-ligand competition assays

Inhibition of CD28–CD80 interaction was evaluated using a competitive ELISA assay with the CD28:B7-1 [Biotinylated] Inhibitor Screening Kit (BPS Bioscience, Cat. #72007), following the manufacturer’s protocol with minor modifications. Briefly, 96-well plates were coated with recombinant human CD28 (2 μg/mL in PBS, 50 μL per well) and incubated overnight at 4 °C. Plates were washed with Immuno Buffer and blocked with Blocking Buffer for 1 h at room temperature.

Serial dilutions of cyclic peptides were first added to CD28-coated wells and incubated for 1 h at room temperature to allow peptide binding. Biotinylated CD80 (5 ng/μL) was then added to the wells and incubated for an additional 1 h to assess inhibition of ligand binding to peptide-occupied CD28. Wells lacking CD28 coating served as ligand controls, while wells treated with inhibitor buffer alone were used as negative controls.

After washing, plates were incubated with streptavidin-HRP (1:1000 dilution in Blocking Buffer) for 1 h at room temperature. Chemiluminescent substrate was added, and luminescence was immediately measured using an Infinite M1000 Pro microplate reader (Tecan, Switzerland). IC_50_ values were determined by fitting concentration-response curves using a four-parameter logistic regression model in GraphPad Prism 10. All experiments were performed in triplicate.

### Cell-based CD28 signaling reporter assays

Functional inhibition of CD28-mediated costimulatory signaling was assessed using the CD28 Blockade Bioassay (Promega, Cat. #JA6101). Jurkat CD28 Effector Cells (2 × 10^4^ cells per well) were seeded in white 96-well plates and pre-incubated with serial dilutions of cyclic peptides (10-point, 1:1 dilution series starting at 200 μM, final 1% DMSO) for 1 hour at room temperature. An anti-CD28 control antibody (Promega, Cat. #K1231) was included as a positive control.

Following peptide pre-incubation, aAPC/Raji cells (2 × 10^4^ cells per well) were added, and co-cultures were incubated for 5 h at 37 °C in a humidified atmosphere containing 5% CO₂. After incubation, Bio-Glo™ Luciferase Reagent (Promega) was added according to the manufacturer’s instructions, and luminescence was measured using a GloMax® Discover System (Promega).

Dose-response curves were generated using GraphPad Prism 10 by fitting data to a four-parameter logistic regression model to determine IC_50_ values. All experiments were performed in triplicate, and data are reported as mean ± SEM.

### Primary human PBMC isolation and culture

PBMCs from healthy donors were obtained from STEMCELL Technologies (Catalog# 70025) and cultured according to manufacturer recommendations. Cells were maintained in RPMI 1640 medium supplemented with 10% fetal bovine serum (FBS), L-glutamine, and penicillin-streptomycin (100 U/mL and 100 µg/mL, respectively) at 37 °C in a humidified atmosphere containing 5% CO_2_. Cell viability following thawing was assessed using trypan blue exclusion prior to experiments.

### T-cell activation assays in primary PBMCs

PBMCs from healthy donors (n = 10 independent donors; STEMCELL Technologies, Catalog# 70025) were cultured in RPMI 1640 supplemented with 10% FBS and antibiotics. Cells were seeded at 2 × 10^5^ cells per well in 96-well plates. T-cell activation was induced using anti-human CD3 (plate-bound, 2 µg/mL) and soluble anti-human CD28 (1 µg/mL). CIP-3 was added at indicated concentrations (0.01-10 µM) at the time of stimulation. FR104 (anti-CD28 monoclonal antibody, MedChemExpress, Catalog# HY-P990587) (10 µg/mL) was included as a benchmark control. Supernatants were collected after 24 h, and IL-2 and IFN-γ concentrations were quantified using ELISA kits (STEMCELL Technologies, Catalog# 02006 and 02003). Each donor sample was tested in technical triplicate.

### Agonist activity assessment

To evaluate intrinsic agonist potential, PBMCs were incubated with cyclic peptides in the absence of stimulation. Cytokine production was measured as described above and compared with baseline controls. Each donor sample was tested in technical triplicate.

### Washout and reversibility experiments

To assess reversibility of CD28 inhibition, primary human PBMCs were preincubated with CIP-3 (1 µM) or FR104 (anti-CD28 monoclonal antibody, MedChemExpress, Catalog# HY-P990587) (10 µg/mL) for 60 min at 37 °C prior to stimulation. Following pretreatment, cells were washed three times with pre-warmed RPMI 1640 medium (centrifugation at 400 × g for 5 min between washes) to remove unbound compound or antibody. Cells were then immediately restimulated with anti-CD3/CD28 activation reagents under the same conditions used in stimulation-only controls. Parallel non-washout conditions were included for comparison.

Cytokine production (IL-2 and IFN-γ) was measured in culture supernatants collected 24 h after restimulation using ELISA (STEMCELL Technologies, Catalog# 02006 and 02003). Functional recovery was calculated as percentage of cytokine production relative to stimulated controls. Experiments were performed using PBMCs from independent donors (n = 10). Statistical comparisons were conducted using one-way ANOVA with multiple comparisons.

### Patient-derived PBMC experiments

PBMCs from patients with ulcerative colitis (n = 5 independent donors; BioIVT, Catalog# HUMANPBMC-0002203) were processed and cultured under identical conditions as healthy donor PBMCs. Cells were obtained from commercial vendors and were provided to investigators in a de-identified manner. No identifiable donor information was available to investigators. Cells were stimulated with anti-CD3/CD28 in the presence of increasing concentrations of CIP-3. Cytokine production was quantified by ELISA after 24 h. Statistical comparisons between peptide and antibody treatment were performed using paired two-tailed t-tests.

### Murine CD28 cross-reactivity assessment

To evaluate functional cross-reactivity of CIP-3 with murine CD28, primary splenocytes were isolated from 8-10-week-old C57BL/6 mice. Spleens were mechanically dissociated through a 70-µm cell strainer to obtain single-cell suspensions. Red blood cells were lysed using ammonium-chloride-potassium (ACK) lysis buffer, and cells were washed twice with RPMI 1640 medium supplemented with 10% FBS, L-glutamine, and penicillin-streptomycin (100 U/mL and 100 µg/mL, respectively).

Splenocytes were seeded at 2 × 10^5^ cells per well in 96-well plates and stimulated with plate-bound anti-mouse CD3 (2 µg/mL) and soluble anti-mouse CD28 (1 µg/mL) antibodies. CIP-3 was added at indicated concentrations (0.01-10 µM) at the time of stimulation. Cells were incubated at 37 °C in 5% CO_2_ for 24 h.

Supernatants were collected and murine IL-2 and IFN-γ concentrations were quantified using ELISA kits (STEMCELL Technologies, Catalog# 02022 and 02020) according to the manufacturer’s instructions. Experiments were performed using splenocytes from independent mice (n = 4 biological replicates), each measured in technical triplicate. Data were analyzed using one-way ANOVA with appropriate post hoc testing.

### Plasma Stability and Microsomal Stability

CIP-3 stability in mouse plasma was evaluated by incubating the peptide (1 µM final concentration) in pooled mouse plasma at 37 °C. Aliquots were collected at 0, 0.5, 1, 2, 4, and 6 h and quenched with ice-cold acetonitrile containing an internal standard to precipitate plasma proteins. Samples were centrifuged at 15,000 × g for 10 min, and supernatants were analyzed by LC-MS/MS. The percentage of remaining parent compound was calculated relative to time zero. Experiments were performed in triplicate.

Metabolic stability was assessed using pooled mouse and human liver microsomes (ThermoFisher Scientific, Catalog# MSMCPL and HMMCPL). CIP-3 (1 µM) was incubated with microsomes (0.5 mg/mL protein) in potassium phosphate buffer (pH 7.4) at 37 °C in the presence of an NADPH-regenerating system. Reactions were initiated by addition of NADPH and terminated at defined time points (0, 5, 15, 30, 45, and 60 min) by addition of ice-cold acetonitrile.

Samples were centrifuged, and supernatants were analyzed by LC-MS/MS. The natural logarithm of percentage remaining compound was plotted versus time to determine the first-order elimination rate constant (k). Intrinsic clearance (Cl_int_) and microsomal half-life (t_1/2_) were calculated using standard equations. Experiments were performed in triplicate.

### Animal studies

Mice were housed and handled according to the guidelines approved by the Institutional Animal Care and Use Committee (IACUC) of our institution (protocol 2023-0028).

### PK analysis of CIP-3 in mice

PK studies were conducted in male C57BL/6 mice (8-10 weeks old; 20–25 g). Animals were housed under standard conditions with ad libitum access to food and water. CIP-3 was formulated in sterile PBS containing 5% DMSO and administered via subcutaneous (s.c.) injection at doses of 1 mg/kg or 5 mg/kg (n = 4 per dose group). Blood samples (∼50-75 µL) were collected via submandibular venipuncture at the following time points post-dose: 0.25, 0.5, 1, 2, 4, 8, 12, and 24 h. Blood was collected into EDTA-coated tubes and centrifuged at 3,000 × g for 10 min at 4 °C to obtain plasma. Plasma samples were stored at −80 °C until analysis.

Plasma concentrations of CIP-3 were quantified using a validated liquid chromatography-tandem mass spectrometry (LC-MS/MS) method. Briefly, plasma proteins were precipitated using acetonitrile containing an internal standard, followed by centrifugation and injection of the supernatant onto a reverse-phase C18 analytical column. Chromatographic separation was achieved using a gradient of water and acetonitrile containing 0.1% formic acid. Detection was performed in positive electrospray ionization mode using multiple reaction monitoring (MRM). Calibration curves were generated using spiked plasma standards over a concentration range of 1-10,000 ng/mL. The lower limit of quantification (LLOQ) was 1 ng/mL. Quality control samples at low, medium, and high concentrations were included in each analytical run to ensure accuracy and precision.

PK parameters, including maximum plasma concentration (C_max_), time to maximum concentration (T_max_), area under the plasma concentration-time curve from 0 to 24 h (AUC_0-24_ h), terminal elimination half-life (t_1/2_), and apparent clearance (CL/F), were calculated using non-compartmental analysis in Phoenix WinNonlin (Certara). Terminal half-life was determined from the slope of the log-linear terminal phase. Data are reported as mean ± SD.

### Adoptive T Cell Transfer Model of Chronic Colitis

Chronic colitis was induced using the adoptive CD4⁺CD45RB^high^ T-cell transfer model as previously described (36–38). Briefly, spleens were harvested from 5-week-old BALB/c donor mice (Jackson Laboratory). Single-cell suspensions were prepared by mechanical dissociation followed by filtration through a 70-µm nylon mesh. CD4⁺ T cells were isolated using magnetic bead separation, and CD4⁺CD45RB^high^ T cells were purified by fluorescence-activated cell sorting (FACS). Sorted cells were washed twice with sterile phosphate-buffered saline (PBS) and resuspended at 1.5 × 10^6^ cells/mL in cold PBS.

Recipient C.B-17 scid mice (7 weeks old; Taconic Biosciences) received 3 × 10^5^ CD4⁺CD45RB^high^ T cells via intravenous injection (200 µL per mouse). Mice were randomly assigned to treatment groups (n = 8 per group) following cell transfer. CIP-3 was formulated in sterile PBS containing 5% DMSO and administered by daily subcutaneous injection at doses of 1 mg/kg or 5 mg/kg. Vehicle-treated mice received formulation buffer alone. Treatment began on day 7 post-transfer and continued for 30 days. Investigators were not blinded to treatment allocation. Disease progression was monitored longitudinally using a disease activity index (DAI) incorporating body weight loss, stool consistency, and fecal blood. Body weight was recorded three times per week. Stool consistency was scored as follows: 0 = normal; 1 = soft but formed; 2 = loose; 3 = diarrhea. Fecal blood was assessed using a guaiac-based test and scored as: 0 = negative; 1 = trace; 2 = positive; 3 = gross bleeding. The DAI score represents the combined average of these parameters.

At study termination (day 30), mice were euthanized and colons were excised, and measured from cecum to rectum under standardized tension. Serum was collected via cardiac puncture and centrifuged at 3,000 × g for 10 min at 4 °C. TNF-α and IL-6 concentrations were quantified using commercially available ELISA kits (ThermoFisher Scientific, Catalog# BMS607-3 and KMC0061) according to the manufacturer’s instructions.

### Statistical analysis

Data are presented as mean ± SEM unless otherwise indicated. Statistical analyses were performed using GraphPad Prism 10.4.1 software. Comparisons between groups were conducted using one-way or two-way analysis of variance (ANOVA) with appropriate post hoc tests as specified in figure legends. *P* values less than 0.05 were considered statistically significant.

## Author contributions

The manuscript was written through contributions of all authors. All authors have given approval to the final version of the manuscript. Conceptualization: M.T.G. and H.D. Methodology: K.K., S.U., and R.Z. Investigation: K.K., S.U., and R.Z. Formal analysis: K.K., S.U., and R.Z. Visualization: K.K., S.U., and R.Z. Supervision: M.T.G. and H.D. Writing-original draft: K.K., S.U., R.Z., M.T.G., and H.D. Writing-review and editing: M.T.G. Funding acquisition: M.T.G. and H.D. Data curation: K.K., S.U., and R.Z. Validation: K.K., S.U., and R.Z. Project administration: M.T.G.

## Competing interests

The authors declare that they have no competing interests.

## Data and materials availability

All data needed to evaluate the conclusions in the paper are present in the paper and/or the Supplementary Materials.

## Notes

### Competing Interest Statement

The authors have declared no competing interest.

## References

1. Pardoll DM. The blockade of immune checkpoints in cancer immunotherapy. Nat Rev Cancer. 2012;12:252–264.

2. Ma K, Xu Y, Cheng H, Tang K, Ma J, Huang B. T cell-based cancer immunotherapy: opportunities and challenges. Science Bulletin. 2025;70(11):1872–90.

3. Wei SC, Duffy CR, Allison JP. Fundamental mechanisms of immune checkpoint blockade therapy. Cancer Discov. 2018;8:1069–1086.

4. Hodi FS et al. Improved survival with ipilimumab. N Engl J Med. 2010;363:711–723.

5. Mehnert JM et al. Safety and antitumor activity of the anti-PD-1 antibody pembrolizumab in patients with advanced, PD-L1-positive papillary or follicular thyroid cancer. BMC Cancer. 2019;19:196.

6. Ribas A, Wolchok JD. Cancer immunotherapy using checkpoint blockade. Science. 2018:359:1350–1355.

7. Su QY, et al. Mechanism and clinical utility of abatacept in the treatment of rheumatoid arthritis. Expert Opin Drug Saf. 2026;25(1):59–70.

8. Upadhyay S, Kaur B, Gabr MT. CD28 and ICOS in immune regulation: Structural insights and therapeutic targeting. Bioorg Med Chem Lett. 2025;127:130310.

9. Bai R, Sun W. Crosstalk between tumor-associated macrophages and the B7/CD28 family in immune checkpoint inhibitor-induced immunotherapy. Mol Cell Biochem. 2026;481(1),127–137.

10. Olejniczak SH, Lotze M T, Skokos D. Second signals for cancer immunotherapy. J Immunother Cancer 2025;13(9):e010530.

11. Lv Y, Jin YL, Zhou Z, et al. The interaction between dendritic cells and T follicular helper cells drives inflammatory bowel disease: a review. Front Immunol. 2026;17:1725349.

12. Ma K, Que W, Hu X, et al. Combinations of anti-GITR antibody and CD28 superagonist ameliorated dextran sodium sulfate-induced mouse colitis. Clin Exp Immunol. 2022;208(3):340–350.

13. Genovese MC et al. Abatacept for rheumatoid arthritis refractory to tumor necrosis factor alpha inhibition. N Engl J Med. 2005;353:1114–1123.

14. Kaplon H, Reichert JM. Antibodies to watch. mAbs. 2021;13:1860476.

15. Leader B, Baca QJ, Golan DE. Protein therapeutics: a summary and pharmacological classification. Nat Rev Drug Discov. 2008;7:21–39.

16. Walsh G. Biopharmaceutical benchmarks 2018. Nat Biotechnol. 2018;36:1136–1145.

17. Scott AM, Wolchok JD, Old LJ. Antibody therapy of cancer. Nat Rev Cancer. 2012;12:278–287.

18. Nelson AL et al. Development trends for human monoclonal antibody therapeutics. Nat Rev Drug Discov. 2010;9:767–774.

19. Chames P et al. Therapeutic antibodies: successes, limitations and hopes for the future. Br J Pharmacol. 2009;157:220–233.

20. Suntharalingam G et al. Cytokine storm in a phase 1 trial of the anti-CD28 monoclonal antibody TGN1412. N Engl J Med. 2006;355:1018–1028.

21. Cierpicki T, Grembecka J. Targeting Protein-Protein Interactions in Hematologic Malignancies. Annu Rev Pathol. 2025;20(1):275–301.

22. Arkin MR, Tang Y, Wells JA. Small-molecule inhibitors of protein-protein interactions: progressing toward the reality. Nat Rev Drug Discov. 2014;21:1102–1114.

23. Strohl WR. Structure and function of therapeutic antibodies approved by the US FDA in 2024. Antib Ther. 2025;8(3):197–237.

24. Ji X, et al. Cyclic Peptides for Drug Development. Angew Chem Int Ed Engl. 2024;63(3):e202308251.

25. Martian, PC et al. Cyclic peptides: A powerful instrument for advancing biomedical nanotechnologies and drug development. J Pharm Biomed Anal. 2024;252:116488.

26. Vinogradov AA, Yin Y, Suga H. Macrocyclic Peptides as Drug Candidates: Recent Progress and Remaining Challenges. J Am Chem Soc. 2019;141:4167–4181.

27. Merz ML, et al. De novo development of small cyclic peptides that are orally bioavailable. Nat Chem Biol. 2024;20(5):624–633.

28. You S, McIntyre G, Passioura T. The coming of age of cyclic peptide drugs: an update on discovery technologies. Expert Opin Drug Discov. 2024;19(8):961–973.

29. Zhang H, Chen S. Cyclic peptide drugs approved in the last two decades (2001-2021). RSC Chem Biol. 2022;3(1):18–31.

30. Jumper J et al. Highly accurate protein structure prediction with AlphaFold. Nature. 2021:596:583–589.

31. Watson JL et al. De novo design of protein structure and function with RFdiffusion. Nature. 2023;620:1089–1100.

32. Hsu C et al. Generative models for protein structures and sequences. Nat Biotechnol. 2024;42:196–199.

33. Stokes JM et al. A Deep Learning Approach to Antibiotic Discovery. Cell. 2020;180:688–702.

34. Lin H. et al. HighPlay: Cyclic Peptide Sequence Design Based on Reinforcement Learning and Protein Structure Prediction. J Med Chem. 2025;68(11):12047–12057.

35. Zhang C. et al. HighFold: accurately predicting structures of cyclic peptides and complexes with head-to-tail and disulfide bridge constraints. Brief Bioinform. 2024;25(3):bbae215.

36. Powrie F et al. Inhibition of Th1 responses prevents inflammatory bowel disease in scid mice reconstituted with CD45RBhi CD4+ T cells. Immunity. 1994;1(7):553–562.

37. Morrissey PJ et al. CD4+ T cells that express high levels of CD45RB induce wasting disease when transferred into congenic severe combined immunodeficient mice. Disease development is prevented by cotransfer of purified CD4+ T cells. J Exp Med. 1993;178(1):237–244.

38. Powrie F et al. A critical role for transforming growth factor-beta but not interleukin 4 in the suppression of T helper type 1-mediated colitis by CD45RB(low) CD4+ T cells. J Exp Med. 1996;183(6):2669–2674.

