## Supporting Information for "AI-designed cyclic peptides enable controllable modulation of the CD28 immune checkpoint"

*Electronic Supplementary Information for*

*Katarzyna Kuncewicz,^a,b#^ Saurabh Upadhyay,^a#^ Renjie Zhu,^c^* *[Hongliang Duan](https://www.sciencedirect.com/author/57213838871/hongliang-duan),^c^ Moustafa T. Gabr^a^**

AUTHOR ADDRESS

^a^Department of Radiology, Molecular Imaging Innovations Institute (MI3), Weill Cornell Medicine, New York, NY 10065, USA

^b^Department of Biomedical Chemistry, Faculty of Chemistry, University of Gdansk, Poland

^c^Faculty of Applied Sciences, Macao Polytechnic University, Macao 999078, China

* To whom correspondence should be addressed:

Moustafa T. Gabr.

^#^These authors contributed equally to this work

| **Contents** |  |
| --- | --- |
| Predicted binding mode of CIP-1 in complex with CD28  Predicted binding mode of CIP-2 in complex with CD28  Mass spectrum of CIP-1 | S3  S4  S5 |
| Mass spectrum of CIP-2 | S5 |
| Mass spectrum of CIP-3 | S6 |
| CIP-3 functionally inhibits murine CD28-mediated T-cell activation. | S7 |

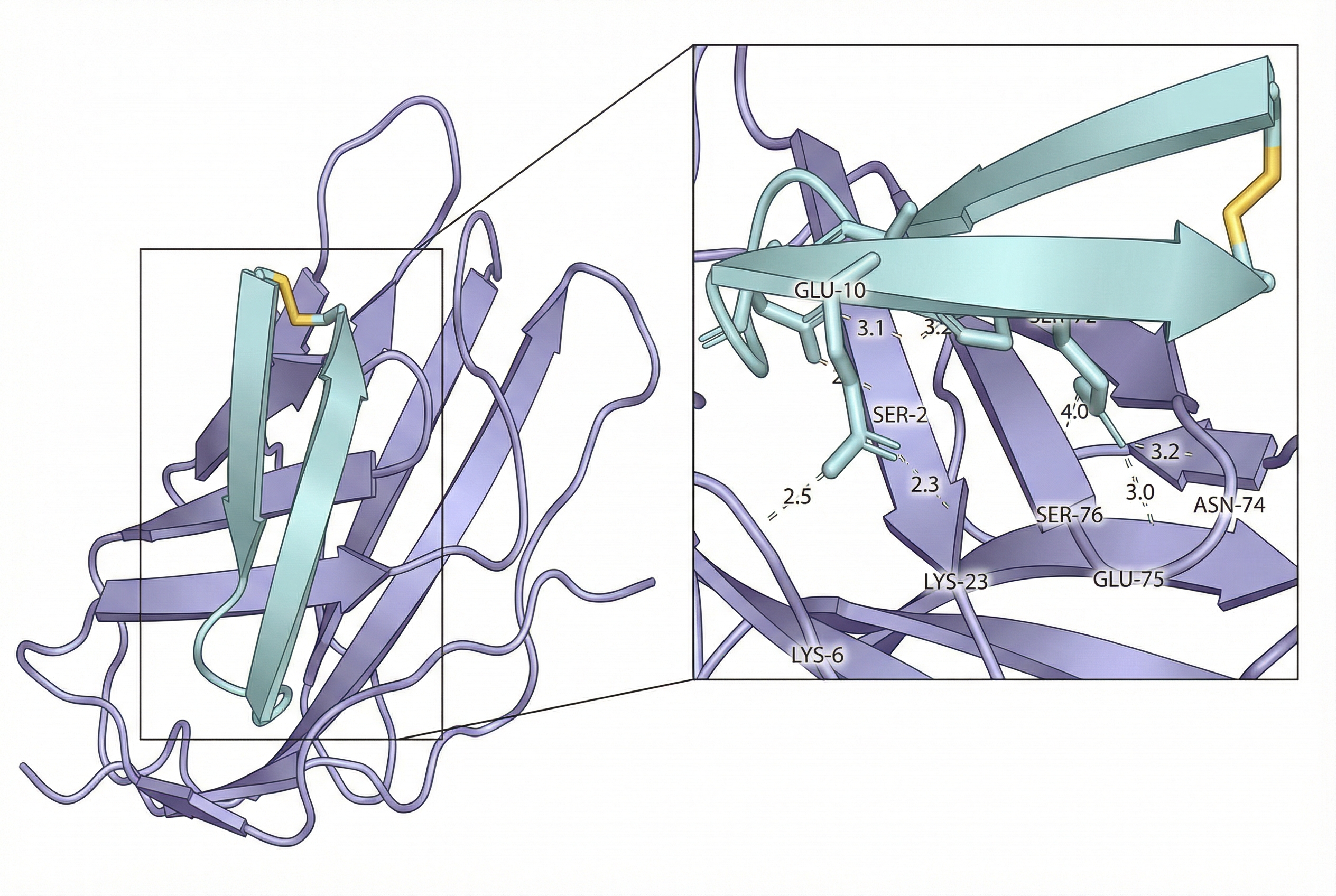

**Figure S1.** Predicted binding mode of CIP-1 in complex with CD28, with inset highlighting key intermolecular interactions and hydrogen bonds stabilizing the interface.

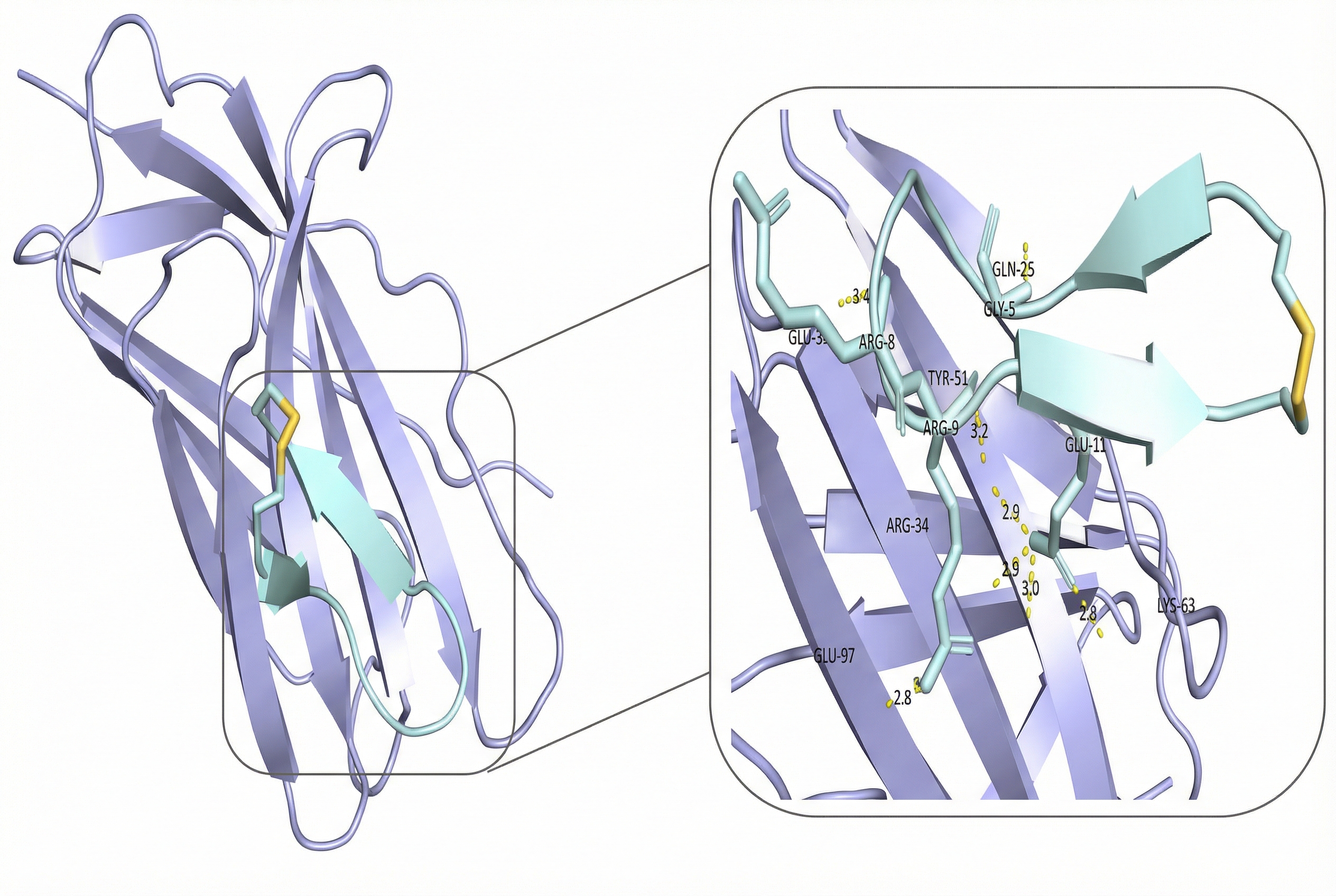

**Figure S2.** Predicted binding mode of CIP-2 in complex with CD28, with inset highlighting key intermolecular interactions and hydrogen bonds stabilizing the interface.

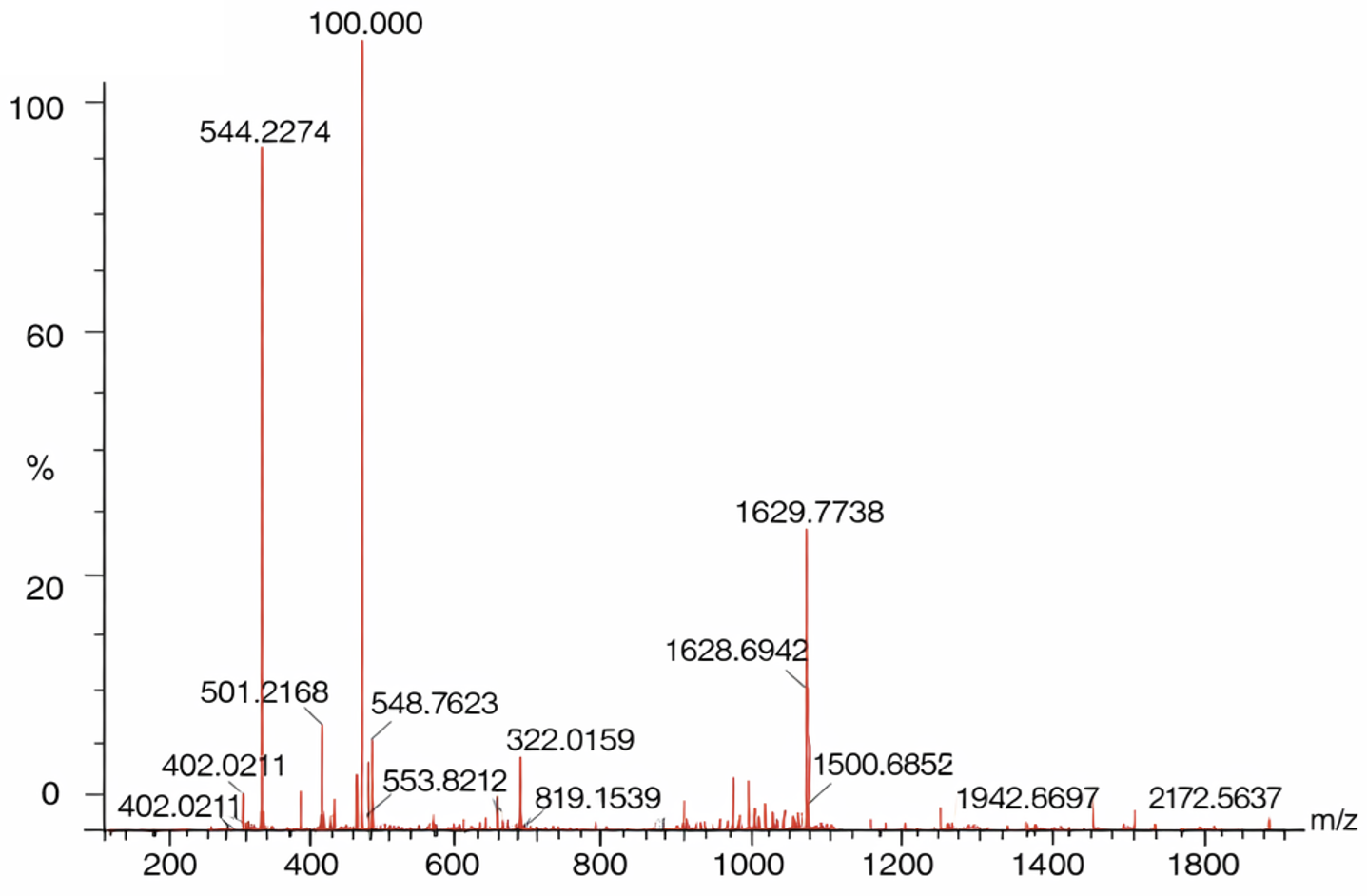

**Figure S3.** Mass spectrum of CIP-1.

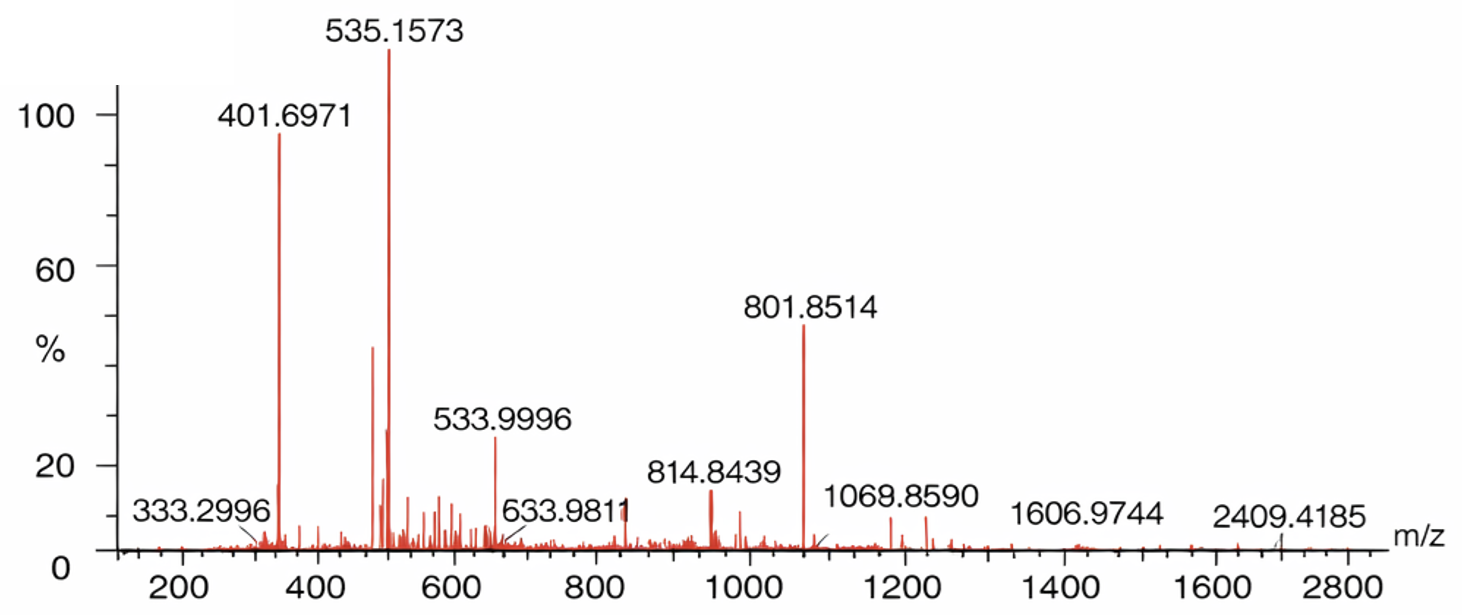

**Figure S4.** Mass spectrum of CIP-2.

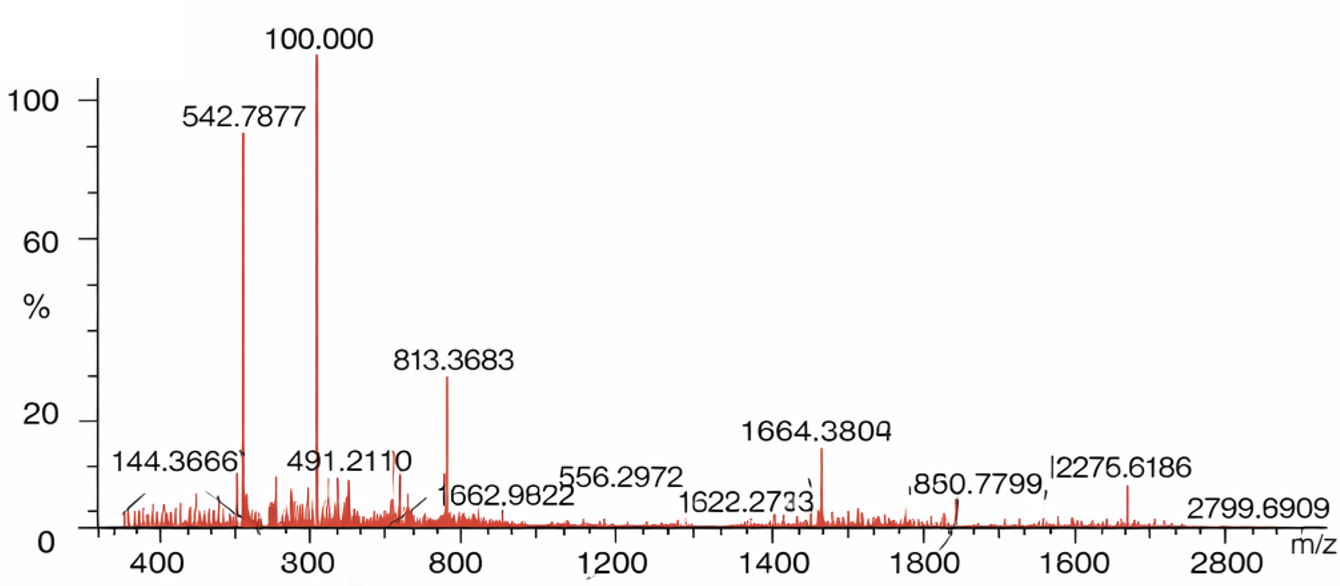

**Figure S5.** Mass spectrum of CIP-3.

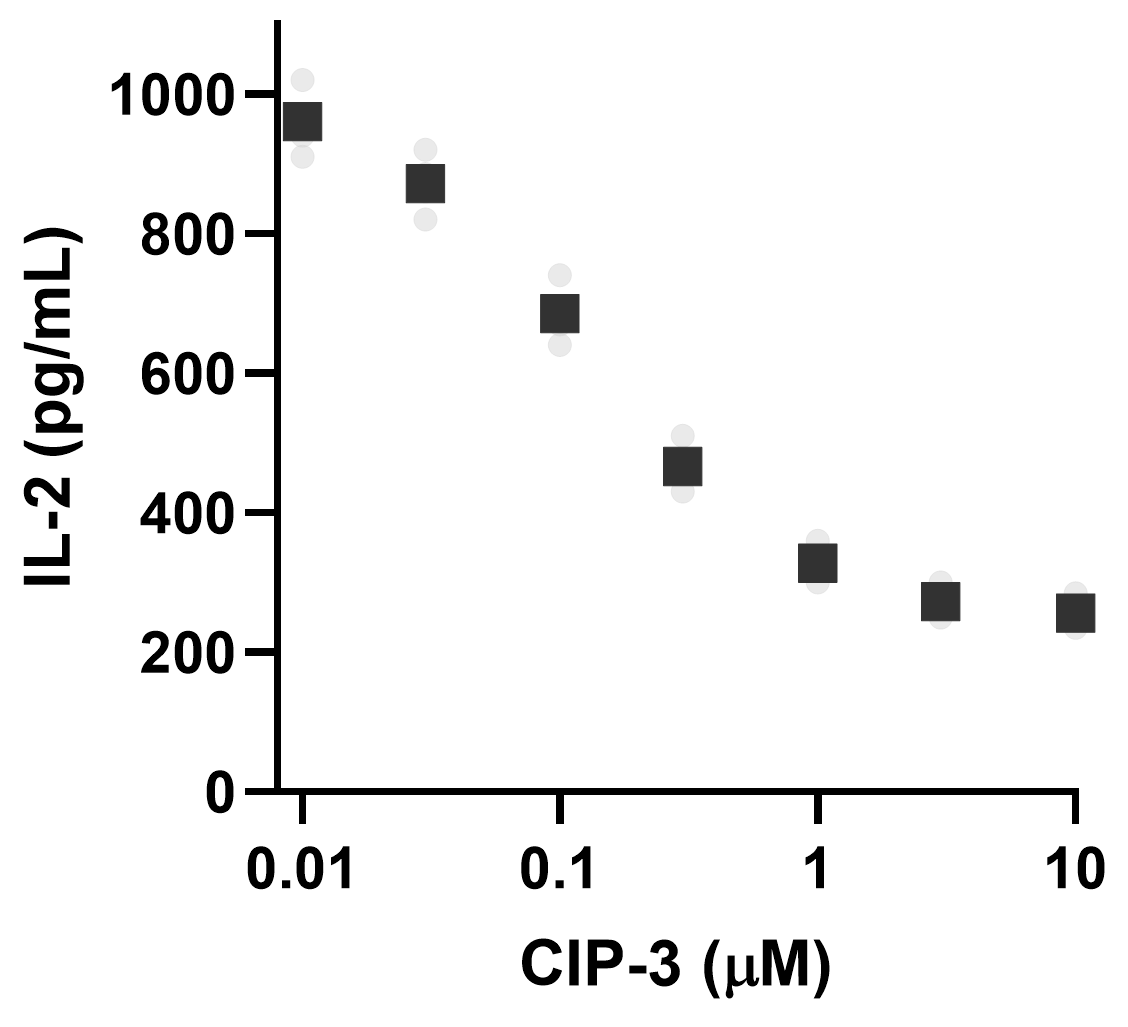

**Figure S6. CIP-3 functionally inhibits murine CD28-mediated T-cell activation.** Primary splenocytes isolated from C57BL/6 mice were stimulated with anti-CD3 and anti-CD28 antibodies in the presence of increasing concentrations of CIP-3 (0.01-10 µM). IL-2 production was quantified after 24 h by ELISA. CIP-3 suppressed murine CD28-dependent T-cell activation in a dose-dependent manner, with an apparent IC_50_ in the submicromolar range comparable to that observed in human PBMC assays. Anti-CD3 stimulation alone produced minimal IL-2 secretion, confirming CD28-dependent costimulatory signaling. Data represent mean ± SEM from independent spleen preparations (n = 3). Statistical analysis was performed using one-way ANOVA with multiple comparisons relative to stimulated control.
